# Data-driven image mechanics (D^2^IM): a deep learning approach to predict displacement and strain fields from undeformed X-ray tomography images - Evaluation of bone mechanics

**DOI:** 10.1101/2023.09.21.558878

**Authors:** Peter Soar, Gianluca Tozzi

**Affiliations:** School of Computing and Mathematical Sciences, Centre for Advanced Simulation and Modelling, Faculty of Engineering and Science, University of Greenwich, United Kingdom; School of Engineering, Centre for Advanced Manufacturing and Materials, Faculty of Engineering and Science, University of Greenwich, United Kingdom

**Keywords:** Bone, X-ray computed tomography, Digital volume correlation, Deep learning, Convolutional neural network

## Abstract

Experimental measurement of displacement and strain fields using techniques such as digital volume correlation (DVC) from in situ X-ray computed tomography (XCT) has notably advanced the understanding of bone mechanics from organ to tissue level. Being experimental in nature, DVC output has been often employed to validate finite element (FE) models of bone improving their predictive ability. Despite the excellent results achieved, these techniques are complex, time consuming, potentially affecting tissue mechanical properties, and their predictive ability requiring prior knowledge of material properties. The recent advent of deep learning (DL) has enabled data-driven models, paving the way for the full exploitation of rich image datasets from which physics can be learnt and retained. Here we propose a novel data-driven image mechanics (D^2^IM) approach based on feed forward convolutional neural network (CNN) that learns from DVC displacement fields of vertebrae, predicting displacement and strain fields for undeformed XCT images. D^2^IM successfully predicted all displacement fields, particularly the one for the z loading axis (*w*), where high correlation (R^2^=0.93) and minimal error (as low as less than 1μm) were found when comparing measured against predicted displacements. The predicted axial strain field in z (*ε_zz_*) was also consistent in distribution with the measured one, displaying generally reduced errors (as low as few tens of με) in the regions within the vertebral body where the effect of border outliers was minimal. This is the first study using experimental full-field measurements on bone structures from DVC to inform DL-based model such as D^2^IM, which represents a major contribution in the prediction of displacement and strain fields only based on the greyscale content of undeformed XCT images. The future development of D^2^IM will incorporate a wider range of structures and loading scenarios for accurate prediction of physical fields in both hard and soft tissues, aiming at clinical translation for improved diagnostics.

## 1. Introduction

The intricate biomechanical properties of musculoskeletal tissues and their response to mechanical loading are crucial for understanding disease effects and optimising treatments. A prime example is bone fracture risk linked to the tissue’s ability to withstand crack propagation, which is closely associated with its distinctive deformation behaviour. Experimental techniques such as digital volume correlation (DVC) are currently considered state-of-the-art in the mechanical characterization of bone, from organ to tissue level. DVC allows for three-dimensional full-field displacement and strain measurements in the material volume by correlating grayscale features from three-dimensional images, typically obtained via high-resolution X-ray computed tomography (XCT) before and after the application of mechanical loading in a process known as in situ XCT mechanics [1]. However, traditional in situ XCT mechanics is performed in a stepwise fashion (i.e. time-lapsed testing), with each loading phase followed by a holding period of 15-30min for full tomographic acquisition to reduce the impact of moving artifacts due to stress relaxation, which is crucial for biological tissues due to their viscoelastic behaviour [2]. Thus, in situ experiments can be considerably time consuming as the XCT imaging (i.e. higher resolution) and mechanical (i.e. increased number of in situ steps) requirements are more demanding, with experiment duration for one specimen close or exceeding the 24h [3, 4]. This can result in extended exposure to X-rays, which is proven to have a substantial impact on the structural and mechanical properties of musculoskeletal tissues both in high-flux synchrotron [5] and laboratory XCT setups [3]. In addition, experimental campaigns to characterise bone using in situ XCT mechanics and DVC are typically conducted with a limited sample size of 2-3 specimens per bone type due to tissue availability, particularly in cases of in vivo treatment [6]. This makes the analysis more qualitative and valid on a case-to-case basis but difficult to generalise within the same cohort, even in the case of tissue samples of similar dimension, extracted from the same anatomical site and tested using the same protocol.

DVC algorithms rely solely on features, for example bone trabeculae, resolved in the acquired XCT images, lacking direct awareness of the object’s material properties. However, the experimental displacement and strain fields from DVC can be used to validate computational models derived from laboratory images, which can then predict mechanical properties under more complex loading scenarios. In this sense, DVC was employed to validate micro-Finite Element (microFE) models, primarily at the biopsy level, focusing on trabecular bone [7, 8]. This validation paradigm has then expanded to encompass more intricate structures, such as vertebral bodies, both with [9] and without lesions [10]. Further developments have seen DVC validation applied to even more complex trabecular bone models featuring heterogeneous material properties [11] and material nonlinearities [12]. Despite the popularity of validation and use of FE models based on DVC, this procedure is technically challenging and based on some material property assumptions varying in accordance with the specific loading regime to simulate (i.e. linear elastic).

The emergence of machine learning (ML), particularly deep learning (DL), has ushered in a new era for the swift resolution of intricate tasks. Different from traditional ML with feature extractors, DL essentially belongs to a class of data-driven end-to-end models, which has achieved great success in different bioengineering areas such as medical image classification [13]. Convolutional neural networks (CNNs) are probably the most popular class of DL models employed in imaging as they possess the ability to learn complex features by extracting visual information automatically using combinations of series of transformations in the model architecture. The typical architecture of CNN has a multi-layer feed-forward network with an input layer, hidden layers including convolutional layers projecting a series of image filers onto the input image and fully connected dense layers, with an output layer where the features will be extracted. CNNs are generally robust with low complexity and easy to train where the network learns throughout the optimization process with a reduced number of parameters [14].

CNNs, have already demonstrated prowess in classifying stages of bone tissue deformation leading to fractures as well as segmenting cracks, employing both high-resolution synchrotron [15, 16] and laboratory XCT [17] in situ mechanics. Recently, DL has been integrated in two-dimensional digital image correlation (DIC) [18–20] and DVC [13]. In both cases, DL has shown remarkable promise by significantly reducing computational complexity, thereby enhancing efficiency in analyses. Additionally, a synergistic approach has been proposed, coupling DL with the traditional cross-correlation method from particle image velocimetry (PIV), to optimize and refine a coarse velocity field, yielding super-resolution calculations [21]. This area holds immense potential for enhancing measurements and advancing models for comprehending and predicting the mechanics of musculoskeletal tissues. Recent advances have seen physics-informed neural networks leveraged to solve problems generally formulated as partial differential equations by predicting full-field data for many processes, notably including crack propagation [22, 23] and mechanical fields such as displacements, stress and strain within a structure [23–27]. In this regard, DL models equipped with the capacity to predict physical fields, such as stress or strain, directly from simple images encapsulating geometry and microstructure information that fully encodes material composition and boundary conditions have emerged [28, 29]. AI-based frameworks have been developed to predict comprehensive strain and stress fields using partial data, enabling the inverse translation from mechanical fields to composite microstructures [30]. However, it’s important to note that both DL-based approaches, whether aimed at enhancing DIC/DVC measurements or predicting physical fields, have primarily been based on synthetic images or patterns and relatively straightforward image geometries and material distributions. Consequently, the potential of these methods to operate on grayscale content and measured full fields from XCT images of complex biological structures, such as bone, remains unexplored.

In the current work, we utilised published datasets of high-resolution in situ XCT images of pristine and artificially lesioned vertebral bone (https://doi.org/10.15131/shef.data.16732441) that were previously used to measure full-field displacement/strain via DVC [31] and validate microFE models [9], simulating the mechanical performance of metastatic vertebrae. We derived a large dataset by augmenting two-dimensional cross-sections of a limited set of three-dimensional XCT tomograms, proposing a new CNN-based approach for data-driven image mechanics (D^2^IM) to predict displacement and consequently strain fields through a synergistic integration of DVC-measured full-field displacements and deep learning. The findings of this study, introduce a major contribution in the prediction of displacement and derived strain fields directly from the grayscale content and texture of undeformed XCT images, setting the scene for advanced image-based prediction of bone deformation and fracture in healthy and pathological conditions.

## 2. Materials and methods

Porcine vertebrae undergoing in situ stepwise XCT compression were used, and a binary mask was made of the unloaded tomography. These were all used as an input for DVC using the python library SPAM to generate full field displacement data. The 3D images of the unloaded tomogram, mask and three displacement fields were all sliced into 2D images, which could be used as an input and ground truth for D^2^IM. D^2^IM is a feed forward Convolutional Neural Network (CNN) that uses four convolutional stacks with max pooling layers. After being flattened, the model feeds into three dense layers which predict the three displacement components. Strains can be derived from both the measured and predicted displacements for comparison. This entire workflow has been summarised in figure 1.

**Figure 1:**
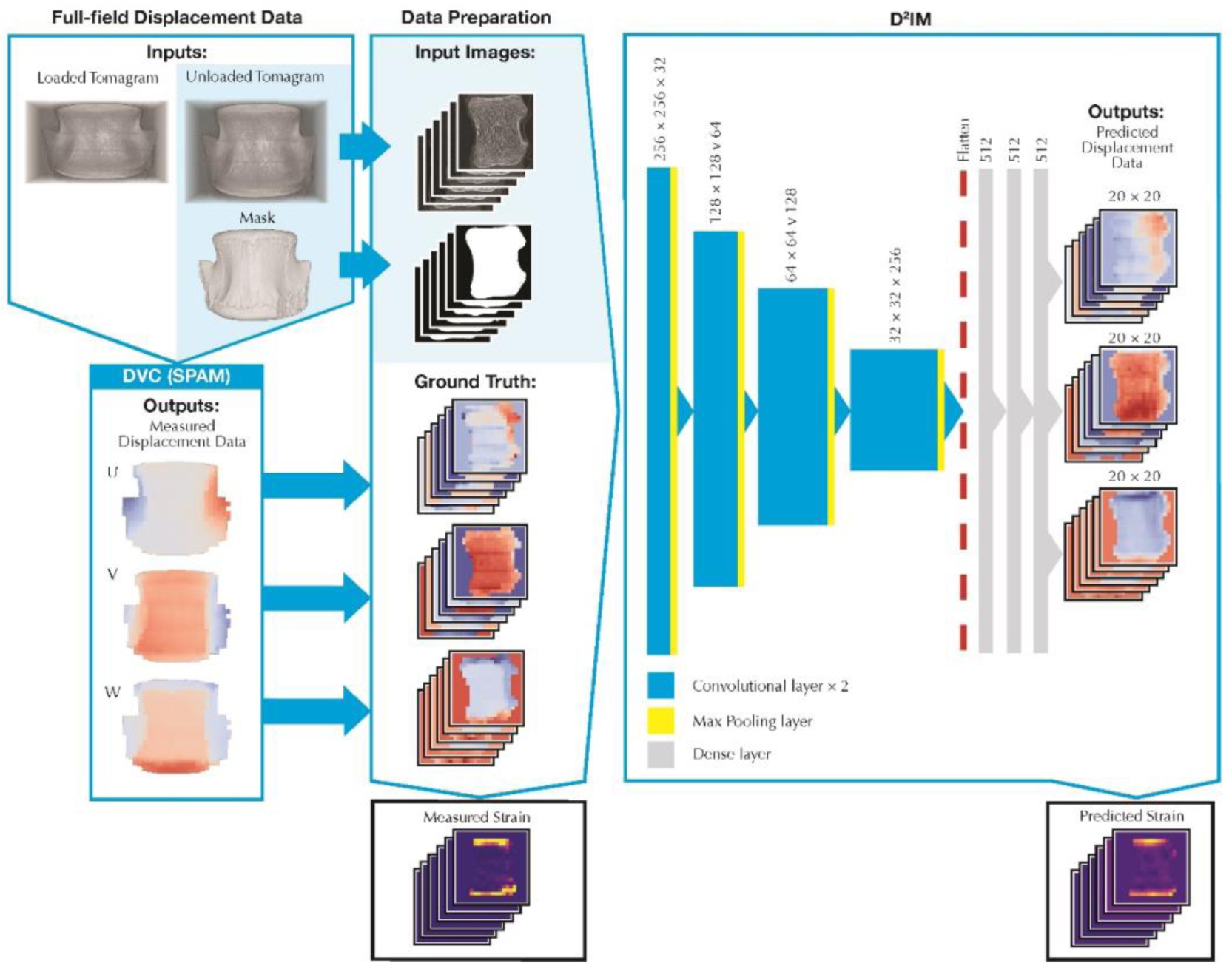
Workflow for generating 2D sliced dataset and making displacement field predictions using Machine Learning Model D^2^IM. Strain fields are then computed from the predicted displacements.

### 2.1 Tomography Dataset

The data chosen to test this modelling framework are a set of XCT images, with a voxel size of 39μm, acquired on ten porcine spine segments with and without artificial lesions prepared and in situ tested by Palanca et al. [9, 31], which have been made available by the authors on Figshare database at https://doi.org/10.15131/shef.data.16732441. This previous work contains the full details of sample preparation, testing and imaging. For each vertebra a tomography has been taken before and after applying a compressive axial load of 6500N. From the unloaded tomography a binary mask (zero outside the vertebral body and one inside) was created for each image using imageJ [32], setting a low threshold value, applying a dilation of 8 voxels, and filling in any remaining internal holes.

### 2.2 Digital Volume Correlation

Digital volume correlation (DVC) was performed with SPAM (Software for Practical Analysis of Materials), an open-source python correlation software operating on a local approach. Further details of the operating principles have already been reported elsewhere [33, 34]. SPAM was used to measure the volumetric displacement fields in vertebrae before and after loading. A non-rigid registration was performed between every loaded and unloaded tomography with a binning of 2 and using the binary mask to identify the region of interest.

The non-rigid registration, raw tomograms and the binary mask are then used in SPAM to measure the full-field displacements in x, y and z; which will henceforth be referred to as u, v and w, respectively. When performing DVC, a window size of 50 voxels was selected, consistently with the optimal nodal spacing of 50 voxels used in BoneDVC software (global approach) by Palanca et al. [9, 31] for the same images. Due to this choice for window size, DVC uncertainties were not computed. However, both displacement and strain were found to be acceptable for vertebral bones with windows/spacing >48 voxels, irrespective of the specific DVC approach [35]. The binary mask was again used to identify the region of interest to measure these displacements, where the correlation window had to contain more than 50% unmasked voxels to be considered an active correlation window by the DVC process. A SPAM filter function was also used on the displacement fields, where the 27 nearest neighbours were considered to remove any outliers.

### 2.3 Machine Learning Dataset Preparation

The proposed machine learning (ML) model required two-dimensional input to output images generated from the three-dimensional unloaded tomography, binary mask and displacement components, which were sliced into sets of 2D images. The image slicing was performed both in the anterior-posterior and medial-lateral direction for each vertebra, as this would serve to both increase numerosity of the dataset and showcase a wider variety of two-dimensional cases.

For the measured displacement components u, v and w all the 50-voxel slices of the 3D images were used, but for their corresponding input data of the unloaded scan and the binary mask only slices corresponding to the central measurement point of each correlation window was kept. All the images have slightly different dimensions and, as the ML model requires consistent image input and output sizes, they had to be resized. The unloaded images from each tomogram were resized to be 256×256 voxels, which is approximately a quarter of the size of the unaltered images, and this resolution was found to be a good compromise allowing for faster training while still maintaining sufficient resolution. The binary mask and three displacement components were all resized to be 20×20 correlation windows, providing a good approximation of the average dimensions of the DVC output across all scans.

To ensure the ML model is learning sensible relationships, sets of images containing an insignificant amount of bone, defined as less than 25% of the image, were removed from the dataset, leaving a final dataset size of 361 image sets. Finally, the unloaded tomography greyscale images were normalised so that their values range between 0 and 1.

### 2.4 Machine Learning Model D^2^IM structure, training and strain prediction

The deep learning (DL) model D^2^IM is a feed forward convolutional neural network (CNN) that takes an unloaded tomography and a binary mask as an input to predict the displacement fields *u*, *v*, and *w* in voxels. The displacement field slices obtained from DVC were used as the ground truth for training and evaluating the accuracy of the model. It should be noted that while all the cases presented in this paper have used the binary mask as an input to improve accuracy of the predictions, the framework can be easily tuned to work without a mask input if required. D^2^IM is able to make a prediction using any greyscale input image, but due to the dataset used for training, it will currently try to interpret greyscale images as vertebral bone structures and predict their deformation under an axial load of 6500N.

The architecture of D^2^IM as feed forward CNN was loosely inspired by VGG16 [36], containing multiple stacked convolutional blocks of increasing depth separated by max pooling layers feeding into multiple dense (fully connected) layers. The key difference comes in the output layer, as rather than being used for the classification of images, D^2^IM is used for regression, predicting the values for the three displacement fields based on the image mappings learnt in the convolutional layers. D^2^IM was created in Python using the machine learning system TensorFlow [37] with a network architecture visually summarised in the right-hand portion of figure 1 and with additional details in table 1. Pairs of convolutional layers are stacked using 3×3 convolutions with zero padding and a stride of 1, followed by a max-pooling layer performed over a 2×2 window with a stride of 2. Four convolutional stacks are used, with the depth of the pairs of convolutional layers progressing until the first stack reaches a kernel size of 32 and each progressive stack doubles in size to achieve kernel size of 256 in stack 4. Once flattened, this is followed by three dense layers of 512 channels, with the third dense layer being fed into the three 20×20 output layers with a multiply function applied to the binary mask that removes any predictions outside the region of interest. These three outputs predict the 20×20 displacement fields *u*, *v*, and *w* for each individual window in the input scan. This model has 36399888 trainable parameters in total, uses ReLU activation functions for every layer (other than outputs having no activation for a regression problem) and uses batch normalisation after every layer. The dense layers are regularised to prevent overfitting using dropout layers, with a rate of 0.5 and L2 regularisation using a factor of 0.00001.

**Table 1:**
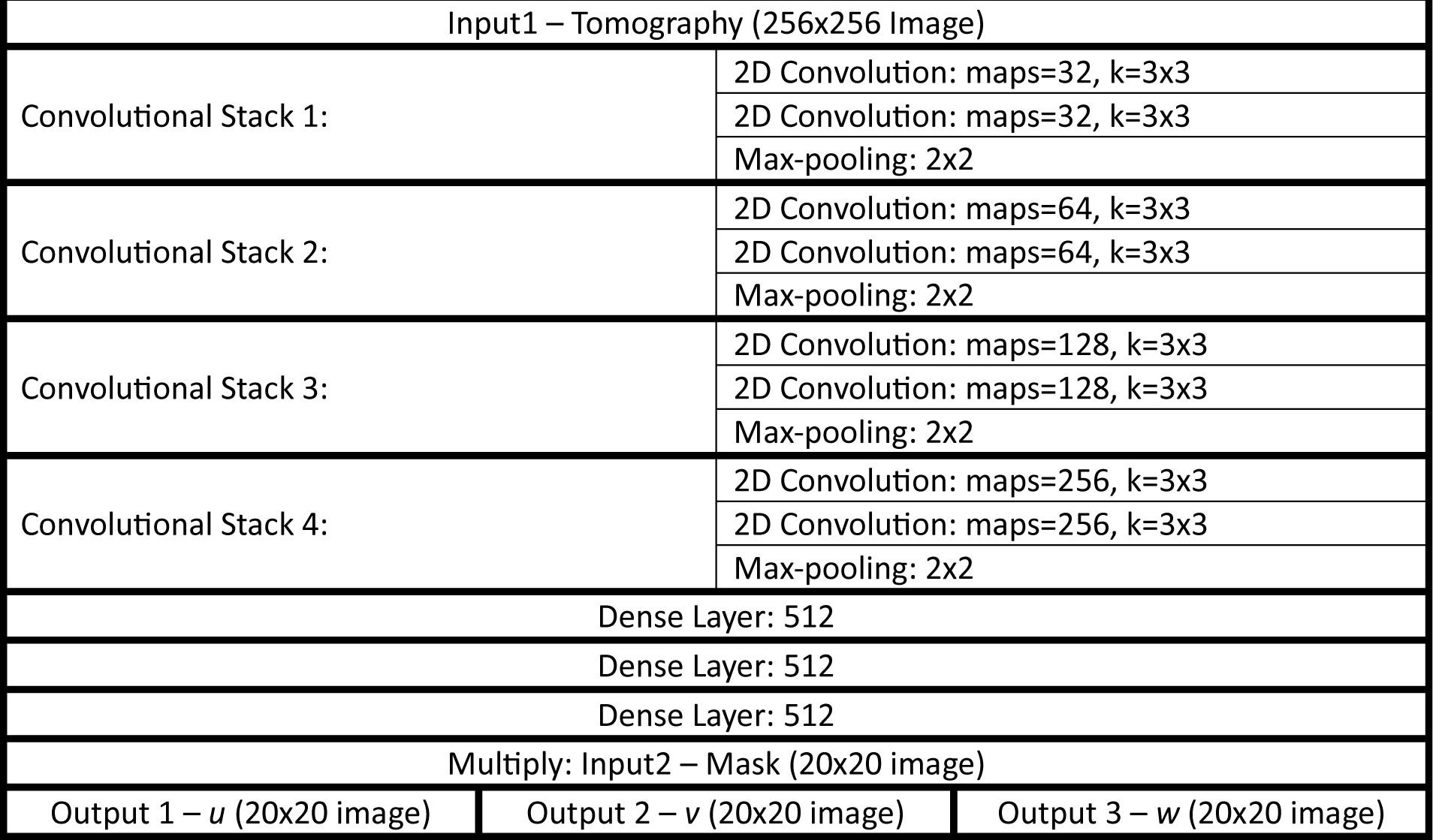
D^2^ IM network Architecture, where ‘maps’ gives the number of output feature maps and ‘k’ provides the convolutional window size.

To train the model, the dataset was shuffled and split into three sets – training (80%), validation (10%) and test (10%), where the training set was used to train the model parameters, the validation set to monitor the model’s performance during training and the test dataset is reserved for the final evaluation of the model’s accuracy. D^2^IM was trained for 500 epochs total with a batch size of 100 using the Adam optimiser to minimise the combined Mean Squared Error (MSE) for predictions of the three displacement components. A learning rate schedule was used, such that the first 300 epochs used a learning rate of 0.001 and the remaining 200 used a learning rate of 0.0001; however, the MSE for the validation dataset seemed to plateau shortly after the 400^th^ epoch.

Strain fields can be derived from both the DVC-measured displacements and predicted using D^2^IM as an alternate way of assessing the performance of the model’s predictions of mechanical behaviour. For this study we focused on comparing measured and predicted normal strains in z (*ε_zz_*), which is the axis of largest displacement due to the experimental setup. To provide consistent visualisation of the results, an additional binary mask (an altered version of the input mask that has been eroded by two pixels) was created to cut off the edge of both strains and errors. This was done for two reasons. Firstly, to exclude unrealistic values of local prediction based on DVC measurement at the borders, which are known to be affected by errors due to progressively reduced amount of greyscale information in those regions [35]. The second reason was to avoid errors between the input mask and the measured DVC output. In DVC, mask information across the correlation window is considered for identifying the region of interest where the displacement should be obtained, but for the input used by D^2^IM only a single mask slice at the centre of the correlation window was used. While this should lead to very similar regions of interest being identified, there are often minor discrepancies at the boundaries of the vertebra, leading to errors being registered where the measured displacement is zero valued but a non-zero value is predicted, or vice versa. Hence, to aid visualisation and provide a more reasonable appraisal of the prediction error, this smaller mask was used such that only cells where the mask input and the ground truth agree were considered.

### 2.5 Error Metrics

Multiple indicators were computed to quantify the errors of predictions made using D^2^IM:

- The performance of the model predictions for each type of displacement field could be individually measured using a Mean Squared Error:

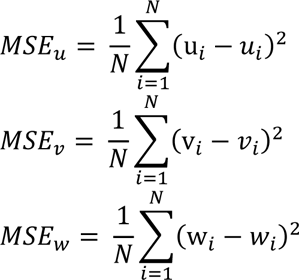

Where N is the number of windows where a displacement value is being predicted; u_*i*_, v_*i*_ and w_*i*_ are the measured displacements in that correlation window; and *u*_*i*_, *v*_*i*_ and *w*_*i*_ are the predicted displacements.

- The overall performance of the model predictions is monitored and evaluated using the sum of the mean squared error of all three displacement fields:

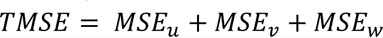

- An absolute error is used for visualization of the error between individual cases of the *w* displacements: |w − *w*| and normal strains in *Z*: |ε_*ZZ*_ − ε_*ZZ*_|.
- The performance of the model predictions of each type of displacement field were also assessed using a relative error as a percentage of the measured value for each predicted window *i*, with these error distributions being used to obtain and visualise the mean and standard deviation of the prediction error.

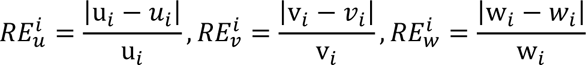

- The variation of the distributions of these relative errors were tested using a Mann Whitney U test at significance level *α* = 0.05.
- The correlation coefficient (*R*^2^) between the measured and predicted displacements were calculated for each displacement component as an alternate means of assessing overall performance in predicting each component.

## 3. Results and Discussion

While both the DVC measurements and D^2^IM predictions represent the displacement in voxels, the results herein presented have been converted to their true displacement values in μm, using the conversion rate of 1 voxel = 39μm as achieved in the original experiment by Palanca et al. [9, 31]. As a first consideration, the DVC approach used for displacement measurement in the current study follows a different correlation strategy (i.e. local approach) than the one in Palanca et al. [31] (i.e. global approach). In brief, local approaches are based on the subdivision of the image into smaller windows (aka sub-volumes) and the spatial correlation of metrics computed in each of the windows of the undeformed and deformed image independently [38]. Global approaches instead are based on the minimisation of the difference of the deformed image and the registered undeformed image when a continuous displacement field is applied [39]. Hence, our first task was to compare displacement magnitude/distribution using SPAM on the same images of Palanca et al. with theirs. As shown in figure 2, the two measurements on the same dataset were consistent indicating how the prediction ability of the proposed D^2^IM approach can be generalised and applicable to any DVC.

**Figure 2:**
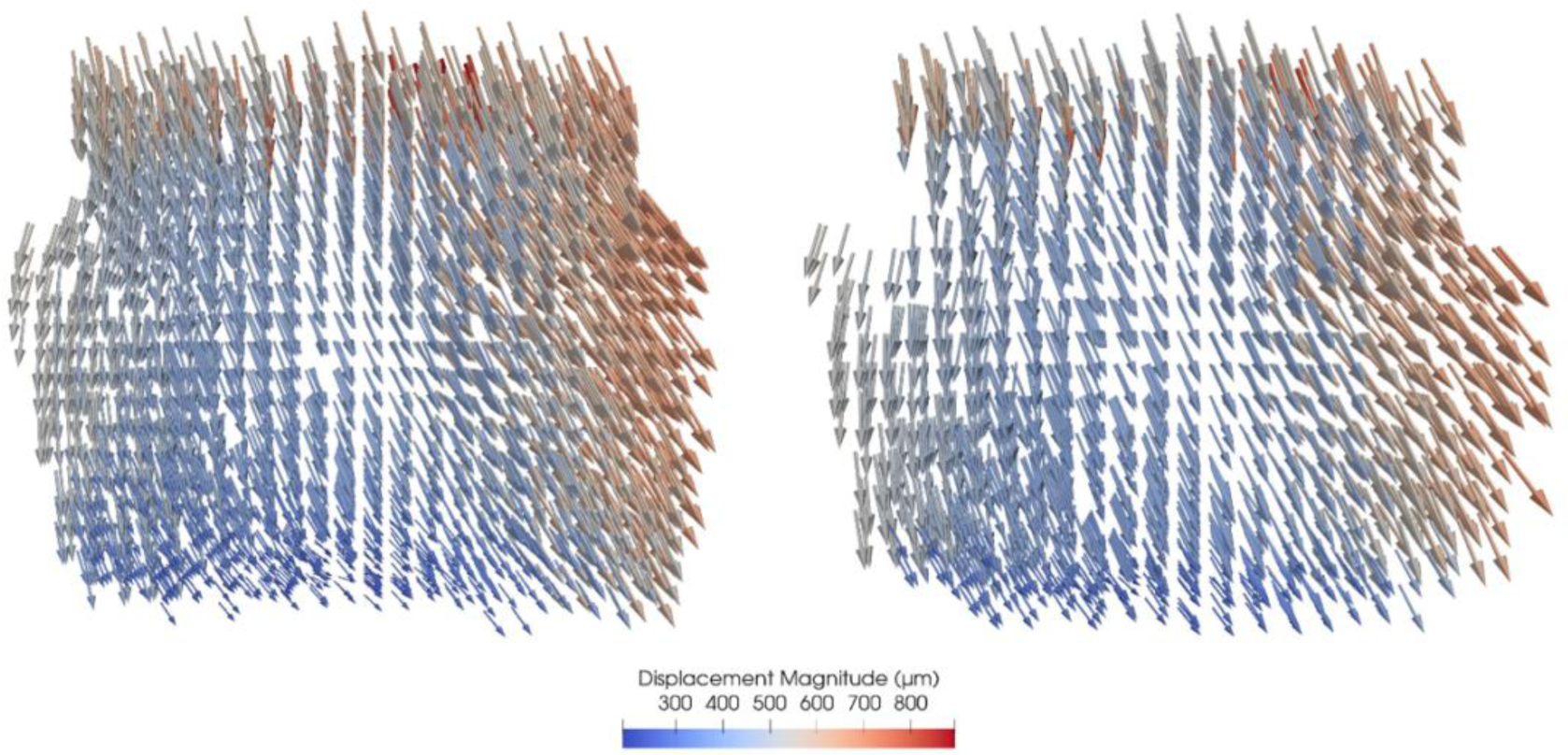
Displacement field magnitude and distribution for in situ XCT of vertebra from Palanca et al [25] (left) and SPAM DVC used in this study on the same dataset (right).

The ability of D^2^IM to generalise across the three displacement components was first examined by making a comparison of the MSE that has been summarised in table 2. Here it can be observed how both validation and test predictions perform similarly with over double the MSE of training predictions for all displacements, showing no indication of data overfitting. On closer inspection, it becomes apparent that the most significant contribution to this deviation in MSE comes from the predictions of the *v* displacement, which was more than doubled in error magnitude compared to the training dataset and largely contributing to the overall error. The test predictions for *u* displacements were also more than doubled in error magnitude compared to the training, but their contribution to the overall error remained relatively minor. The *w* displacements demonstrated the most consistent prediction behaviour by this metric and slightly increasing MSE in the validation and test predictions when compared to training.

**Table 2:**
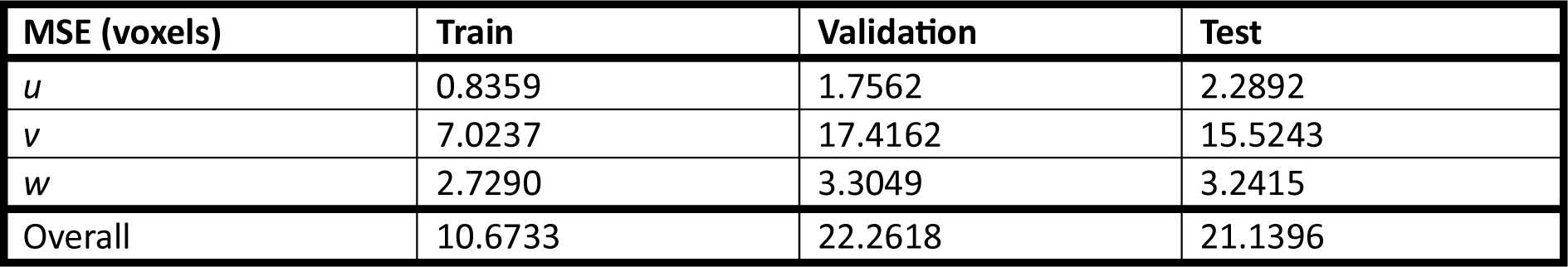
MSE for D IM displacement predictions in voxels for the training, validation, and test data after training for 500 epochs.

However, the raw MSE overlooks the fact that data measured on a larger scale will likely have an intrinsically higher MSE regardless of the prediction accuracy, hence summary statistics describing the distribution of displacement values across every window were obtained for the measured and predicted displacements in the test dataset, as summarised in table 3.

**Table 3:**
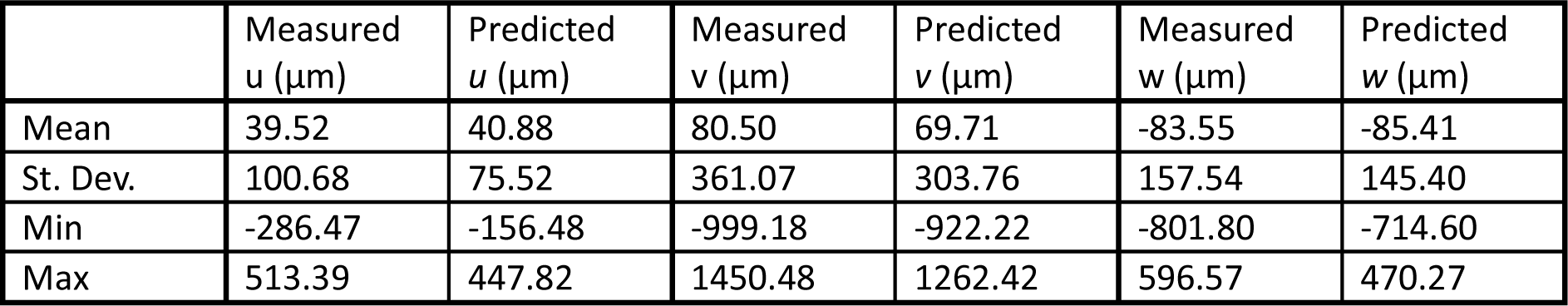
Summary statistics for the measured and predicted displacement fields of the test dataset.

Overall, these statistics seem to demonstrate how the fundamental behaviour is being captured for all three displacement components, with none of the statistics for the predicted values being remarkably different to those measured. At the same time, it does also reinforce the overall trends observed in the MSE, with the *u* displacement’s relatively small MSE being partially explained with a mean value half the size of the other two components in absolute terms. The *v* displacements have by far the largest range in measured values, which could relate to the large MSE observed even for the training predictions (table 2). The poor behaviour in the test predictions for *v* can also be observed with the statistics for such values having some of the biggest discrepancies from measurements, most notably in the mean value that was significantly lower for the predicted values than the measured ones. Finally, the *w* displacements continued to show the most consistent prediction behaviour based on the mean and standard deviation, with some discrepancies between the predicted and measured minimum and maximum values, which is consistent with a typical trend of under-predicting high values and over-predicting low values already reported in literature [28, 40].

A correlation analysis (figure 3) was then performed between all the measured and predicted displacements to further explore the error in the predictions. This again highlighted a very good accuracy of the predictions, with strong positive correlation coefficients for all three displacement components. While there are clear outliers and scatterings of poor predictions, they mostly make up a relatively minor proportion of the 7584 non-masked windows being predicted and plotted. The *u* displacements had the lowest R^2^ value of 0.85, with the most egregious scattering of poor predictions, especially for the more extreme valued displacements. While the *v* displacements reported the largest R^2^ value of 0.95, inspecting the scatter charts the *w* displacements seemed to produce the most consistent behaviour as the *v* displacements displayed some clear clustering of poor predictions, along with more significant outliers.

**Figure 3:**
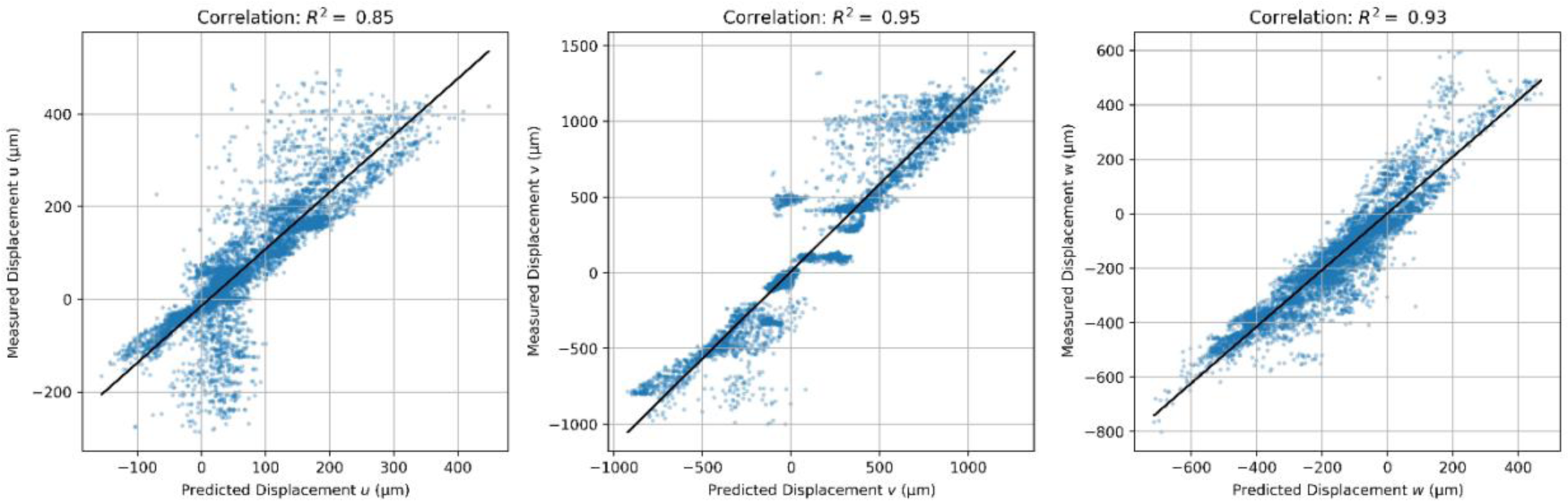
Correlation analysis between all measured and predicted displacements for the test dataset, performed across all three displacement components, including a line of best fit and the calculated correlation coefficient R^2^.

The relative error of the predictions was also examined and reported in figure 4, where the error of a predicted displacement is represented as a percentage of the measured value in the window.

**Figure 4:**
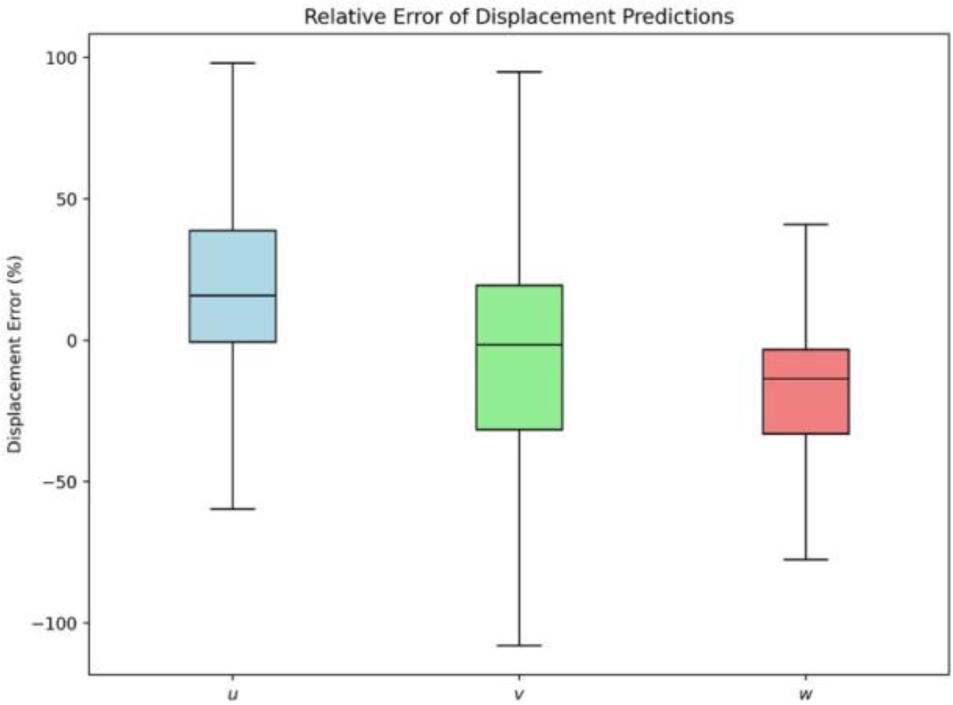
Box and whisker chart with relative error distribution as a percentage of the measured value across the three displacement components. Boxes show the interquartile range and whiskers the maximum and minimum observed errors (excluding outliers).

This reinforced previous findings as most predictions for all three displacements were fairly accurate, with the relative displacement errors having median values close to zero and interquartile ranges of 37% for *u*, 51% for *v* and 27% for *w*. Once again, while some large errors were observed for the v displacements and these were considerably reduced for w (p < 0.05), which continued to show the most consistent behaviour with the tightest error distribution.

While all displacement components predicted by D^2^IM seemed to capture the overall relationships with a reasonable degree of accuracy, these metrics consistently showed the *w* displacements in z as the best performing component. This is encouraging as the framework implemented in D^2^IM for this study mainly aimed at predictions based on bone loaded in compression, where the largest displacements are expected in z and normal strain fields in that direction (ε*_zz_*) are prevalently used in describing the local mechanics [41, 42] The ability of D^2^IM in predicting full-field displacements (*w*) and strains (*ε_zz_*) is reported for pristine and lesioned vertebral sections sliced in both anterior-posterior and lateral-lateral directions (Figure 5-8).

**Figure 5:**
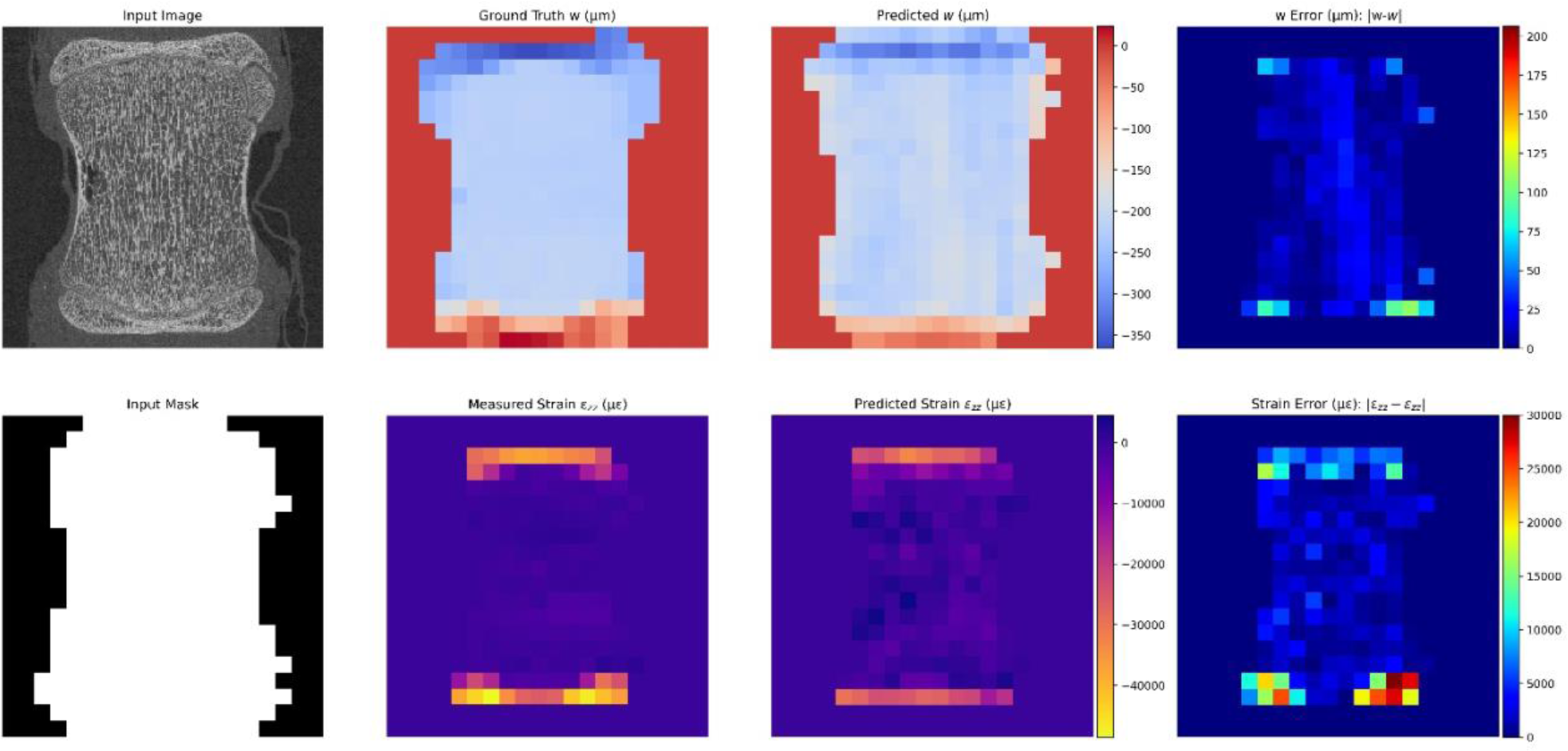
Results for an anterior-posterior sliced tomography with no lesion. Top row left-right: Input tomography slice, measured displacements, predicted displacements and absolute error of displacements. Bottom row left-right: Input binary mask slice, measured strain, predicted strain and absolute error of strain.

The first image is of a pristine anterior-posterior sliced vertebra as shown in figure 5.

D^2^IM is predicting very similar behaviour for both displacement and strain fields to those measured using DVC. The displacements both showed the largest magnitudes (∼366μm for w and ∼337μm for *w*) at the top endplate and a relatively consistent displacement throughout most of the vertebral body, before rapidly shrinking to nearly zero towards the bottom endplate. This was matched by both strain profiles having a bar of similar high magnitude strain at the top and bottom endplates (∼48532με for ε*_zz_*and ∼33082με for *ε_zz_*), with comparatively small strain being registered throughout the rest of the structure. Errors were generally small and without much noticeable pattern within the vertebral body (as low as 0.04μm for displacement and 26με for strain), except at the bottom of the structure where there is a mismatch in the displacement and strain behaviour. Some outliers were localised at the borders/corner of the masked image and, although the second masking strategy was successfully implemented in D^2^IM, there may be instances in the visualisation with errors still caused by either DVC boundary uncertainties in the measurement or mismatch occurring within the smaller mask.

The second case is on another anterior-posterior sliced image from a vertebra with artificial lesion (figure 6).

**Figure 6:**
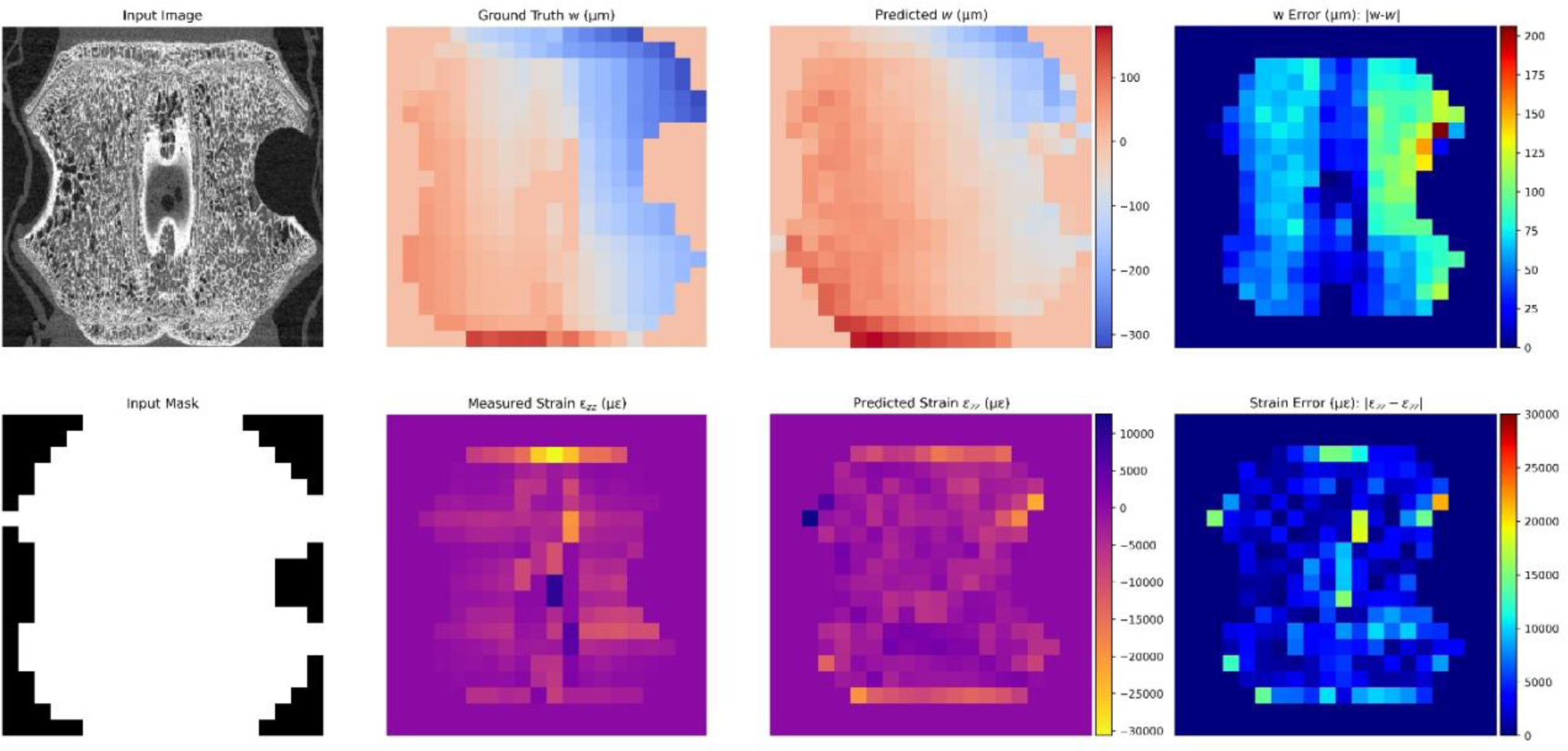
Results for an anterior-posterior sliced tomography with lesion. Top row left-right: Input tomography slice, measured displacements, predicted displacements and absolute error of displacements. Bottom row left-right: Input binary mask slice, measured strain, predicted strain and absolute error of strain.

D^2^IM captured the underlying displacement and strain behaviour, with the lesioned side of the structure experiencing negative displacements in z, while the non-lesioned side showed a positive displacement. The strain was quite small for both the measured and predicted output, barring a line of high strain magnitude (∼30661με for ε*_zz_*and ∼19655με for *ε_zz_*) at the top endplate and some outliers visibly caused in this case by a masking mismatch. While the underlying behaviour is being captured, there was more of a systematic error when compared to the pristine case in figure 5. This is believed primarily to be an issue of numerosity, as while an equal number of five cases with and without lesions were used, the position of the lesion in the vertebra differed between the five lesioned cases. Specifically, three of the lesions were in the anterior position of the vertebra and only two had a lesion in a lateral position, which is the scenario presented here. Consequently, due to having three less directly applicable cases to learn from, it is intuitive that D^2^IM found the lateral lesion scenario to be more challenging to predict.

In figure 7 an image of a pristine vertebra sliced in the lateral-lateral direction is presented. Similarly to the pristine anterior-posterior case, there was a very good agreement between prediction and measurement. Both identified a band of large negative displacement at the top endplate of the vertebra (maximum ∼802μm for w and ∼726μm for *w*) and running down the right side for the top half of the body. Most of the structure showed relatively consistent smaller displacements, before reaching a band of nearly zero displacements along the bottom endplate of the vertebra. It must be noted that while D^2^IM identified the top right being the point of highest displacement magnitude with a band of large displacements going down the right edge, the predictions were smaller in magnitude than the measured. This error was carried across into the strains, where both the predicted and measured strain fields showed similar bands of strain at the top and bottom endplate, but the predictions did not capture the vertical band of strain going down the top-right edge of the vertebra.

**Figure 7:**
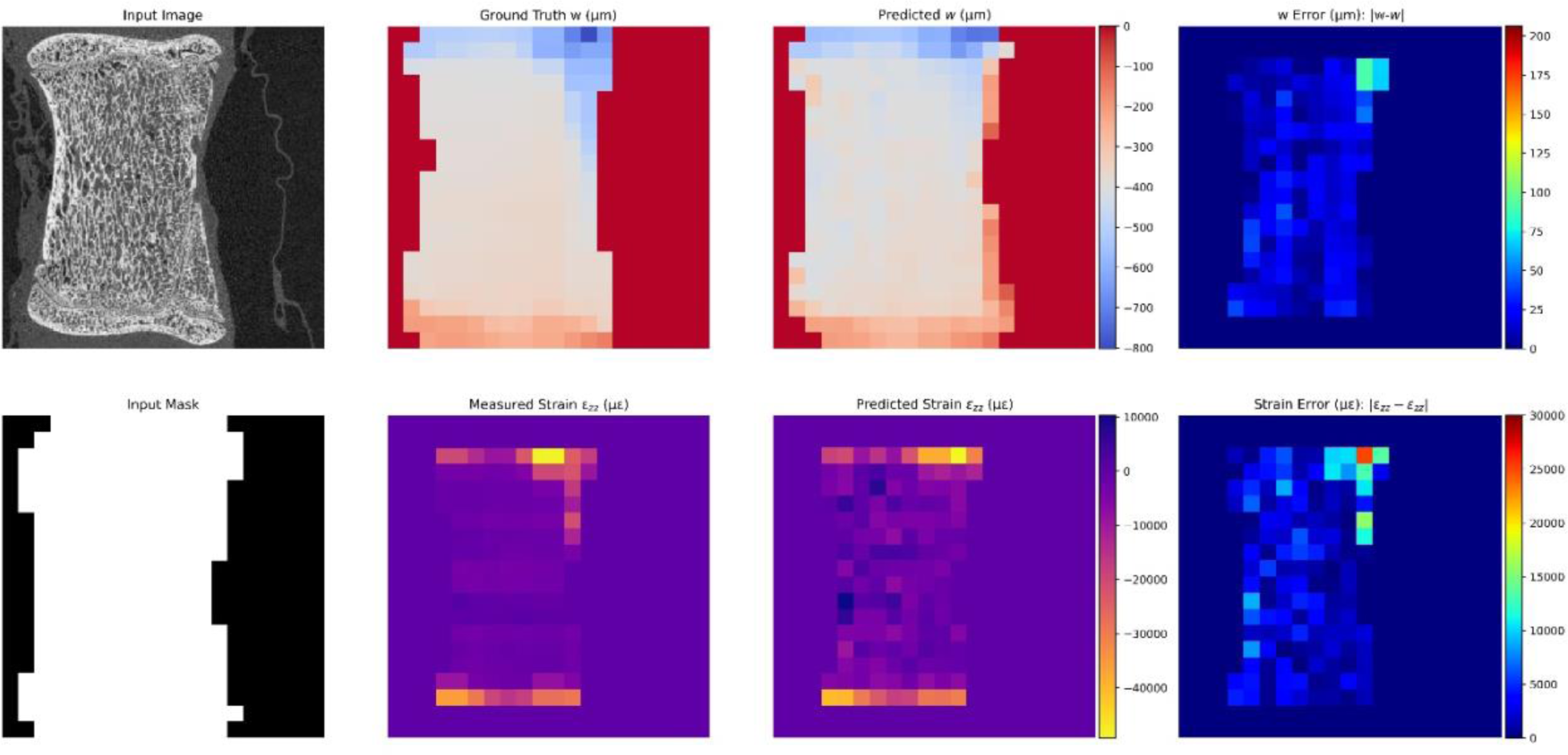
Results for a lateral-lateral sliced tomography with no lesion. Top row left-right: Input tomography slice, measured displacements, predicted displacements and absolute error of displacements. Bottom row left-right: Input binary mask slice, measured strain, predicted strain and absolute error of strain.

The final case being presented (figure 8) is on the prediction results for a vertebra with a lesion and sliced in a lateral-lateral direction.

**Figure 8:**
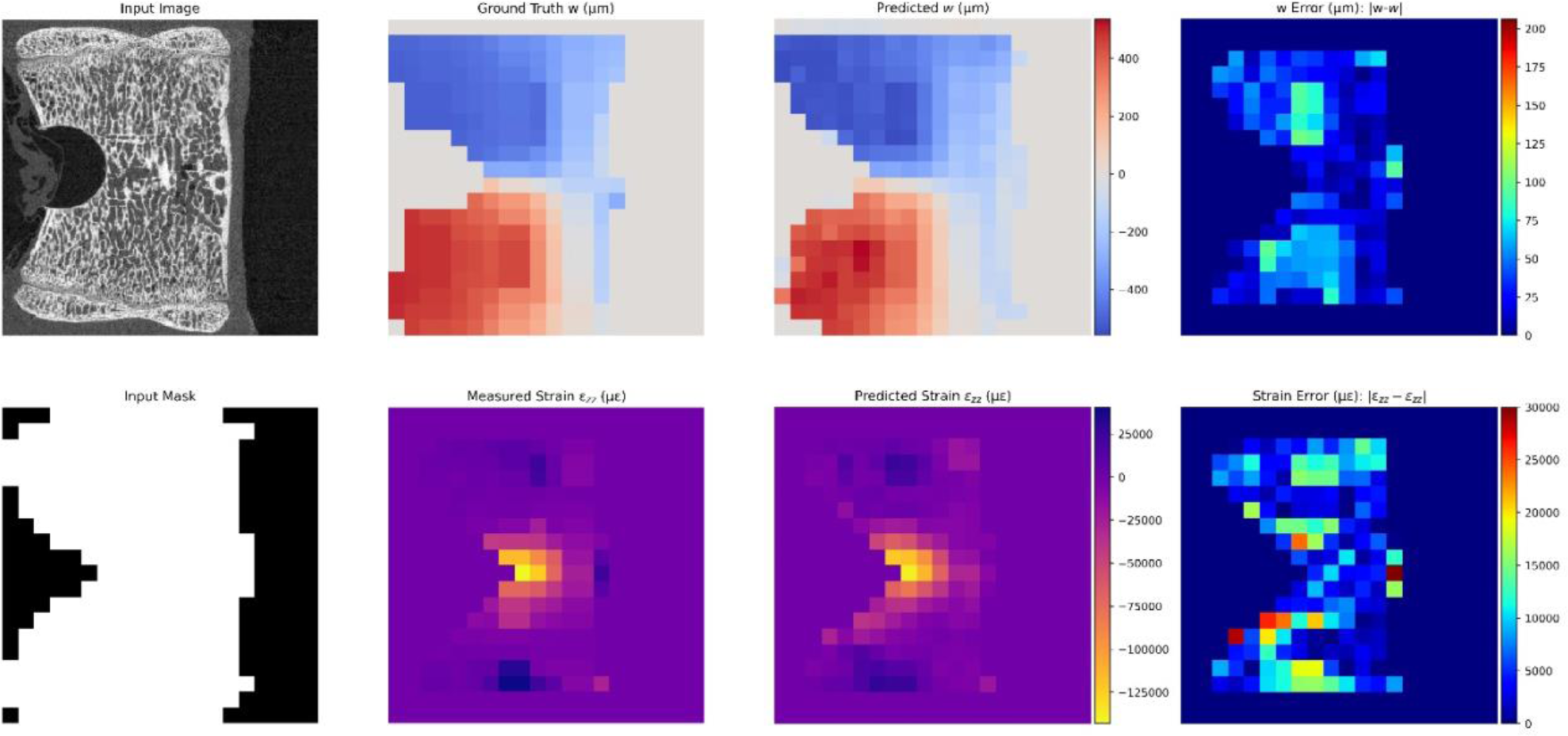
Results for a lateral-lateral sliced tomography with lesion. Top row left-right: Input tomography slice, measured displacements, predicted displacements and absolute error of displacements. Bottom row left-right: Input binary mask slice, measured strain, predicted strain and absolute error of strain.

The results showed very good agreement in the general behaviour, with less of a notable systematic error than in the other case presented with a lesion in figure 6, likely due to the presence of an additional case with an anterior lesion in the dataset. While the displacements were generally slightly smaller in magnitude for the predictions, the underlying behaviour is clearly being caught with negative displacements above the lesion and positive displacements below, both increasing in magnitude as they move towards the front of the vertebra. The displacement errors are generally small and seemingly random, barring two square regions above and below the lesion where more significant errors are observed (up to 101μm), which was perhaps also an issue related to less available cases with an anterior lesion than of pristine vertebra. The strains reported a similar good match between the predicted and measured values, with both cases showing a large negative strain near the lesion in the centre of the vertebra (∼143179με for ε*_zz_* and ∼138375με for *ε_zz_*), with some smaller, but still significant positive strains near the top and bottom endplates. The error in strain is generally small, barring border outliers as in previous cases, without a noticeable pattern.

Based on the initial rationale on which D^2^IM was developed, the superior performance of the *w* displacements was expected when considering the true nature of the problem and the simplification of using 2D slices. The real experiment is of course a 3D problem, with the entire morphology of the structure playing a role in the structural behaviour, leading to the displacement components measured via DVC. Hence, when taking a slice of the structure, a great deal of greyscale information required to make an accurate prediction of the displacements is not considered. For example, in this dataset half of the data was from vertebrae with an artificial lesion, which impacted on the observed structural behaviour but, as evidence of this lesion is generally not found in all slices, D^2^IM may be presented with two almost identical slices of which one was from a lesioned vertebra and the other was not. This means that the measured displacements could differ for these slices, as there is no additional information in the input images that D^2^IM can learn to account for this. While this could represent a potential problem for all displacement components, the *u* and *v* displacements were most affected. In fact, depending on the slicing direction (note that anterior-posterior was slicing in y and lateral-lateral slicing in x for our coordinate system), one of these components will always be deforming into the slicing direction, often meaning sufficient information is lacking to make an accurate prediction. However, while missing information outside the slices represented an issue in some cases, the *w* displacements experienced this to a much less significant degree as all images in the dataset included the caudal-cranial (z) change in structure, whichever direction slicing was performed. Hence, the input structure was always axial to the force being applied, meaning the slices generally contained enough structural information on the observed displacements to make a reasonable prediction.

The strain fields predicted by D^2^IM were in good agreement with those measured by DVC. For this study we focused on comparing the normal strains in z, which is on the axis of largest displacement due to the experimental setup. Due to the combination of anterior-posterior and lateral-lateral slices of the vertebrae being used in the dataset, calculation of the other normal strains, shear strains or any of the principal strains were excluded. If included there would be a requirement to identify the type of slicing used for each image before these other strain elements could be correctly calculated. Furthermore, while all three displacements are being predicted, they were represented on a 2D slice, so constructing 3D strains would be problematic and require information going into the page of the slice that is not easily estimated or obtained with the current outputs of D^2^IM. In any case, and despite only one strain output presented, the local mechanics predicted by D^2^IM is in line with that presented other studies using DVC on pristine vertebrae [31, 43], where high compressive strain localised in correspondence to endplates and reaching magnitudes up to 55000 microstrain. In the presence of lesions. Palanca et al. reported high localised strains, with magnitudes in the range 80000-200000 microstrain [9], which is consistent with D^2^IM prediction. A further consideration is needed for predicted strain fields. Given the strategy used to increase numerosity with 2D XCT slices and the conscious choice of structuring D^2^IM to use the primary DVC output, strains are being derived directly from predicted displacement fields and not ‘learned’. Thus, they carry across the displacement errors, mainly resulting in some underestimation of higher strain magnitudes and some overestimation of lower strain magnitudes (figures 5 and 7). Whilst the former remains representative of regions of tissue already exceeding yielding (i.e. 10000με for bone in compression [44]), the latter could indicate a prediction of yielded portions of the tissue that weren’t measured as such, as shown in figure 9.

**Figure 9:**
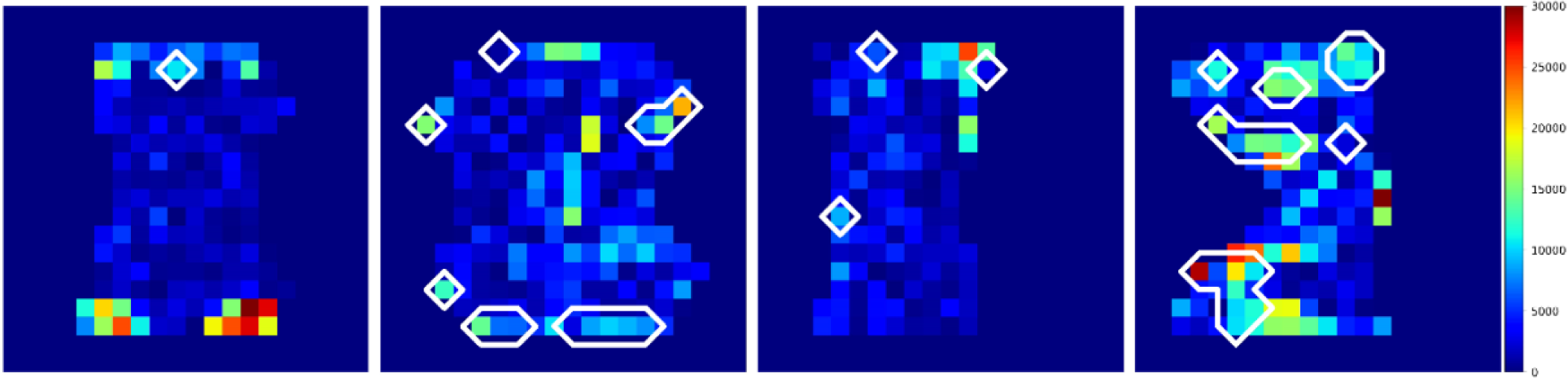
Bone yield strain error for the different cases reported in figures 5-8 (left-right). The marked areas correspond to measured strain in DVC windows that was lower than 10000με, whereas predicted strain exceeded 10000με.

It was found that incongruences in strain predictions exceeding the set threshold of 10000με, when compared to measured values which did not, were mostly localised in border outliers and in proximity of regions where bone was either yielded or pre-yielding. This is particularly evident in the last case from figure 8 (far right in figure 9), reporting the largest range of displacement and higher strain magnitude close to the lesion where errors are smaller. Once again, such discrepancies may be attributed to the reduced number of lesioned vertebrae in a specific location, as well as strain values an order of magnitude higher than tissue yielding which could be misinterpreted by D^2^IM at this stage of development. However, it must be noted that the above-mentioned singularities had a very limited incidence on the overall distribution and mechanical relevance of the predictions for all cases.

## 4. Conclusion

In this study a novel data-driven image mechanics (D^2^IM) approach was developed to predict displacement and strain fields directly from undeformed XCT images. D^2^IM learning used experimentally measured displacement fields for vertebrae, with and without artificial lesions, obtained via digital volume correlation (DVC). D^2^IM successfully predicted all displacement fields and displayed an excellent performance for both displacement and strain fields in the main loading direction of the vertebra. The findings represent a breakthrough in the prediction of physical fields by exploiting only the rich information contained in undeformed XCT images. The future development of D^2^IM will aim at learning a wider range of mechanical data from both hard and soft tissue in healthy and pathological conditions for accurate prediction of physical fields, avoiding long, repetitive and tissue-damaging experimental campaigns but also to advance integration of data in more sophisticated FE simulations. Ultimately, translating into clinical imaging to develop the next generation of diagnostic techniques.

## Data availability statement

D^2^IM code and derived data supporting the findings of this study are available on request from the corresponding author.

## Author statement

### Peter Soar and Gianluca Tozzi

Conceptualization of the methodology, data analysis, and drafting, editing, reviewing and finalizing the paper. **Peter Soar**: Digital volume correlation analysis, software/code development and execution, modelling, data collection and visualisation.

## Acknowledgements

The authors acknowledge support from the School of Engineering and the School of Computing and Mathematical Sciences at the Faculty of Engineering and Science.

